# Emerging variants of canine enteric coronavirus associated with seasonal outbreaks of severe canine gastroenteric disease

**DOI:** 10.1101/2022.10.03.510536

**Authors:** Edward Cunningham-Oakes, Jack Pilgrim, Alistair C. Darby, Charlotte Appleton, Chris Jewell, Barry Rowlingson, Carmen Tamayo Cuartero, Richard Newton, Fernando Sánchez-Vizcaíno, Ivo Salgueiro Fins, Bethaney Brant, Shirley Smith, Rebekah Penrice-Randal, Simon R. Clegg, Ashley P.E. Roberts, Stefan H. Millson, Gina L. Pinchbeck, P-J. M. Noble, Alan D. Radford

## Abstract

Canine enteric coronavirus (CECoV) variants have an emerging role in severe outbreaks of canine gastroenteritis. Here we used syndromic health data from a sentinel network of UK veterinary practices to identify an outbreak of severe canine gastroenteritis. Affected dogs frequently presented with vomiting, diarrhoea and inappetence. Data from sentinel diagnostic laboratories showed similar seasonal increases in CECoV diagnosis. Membrane glycoprotein (M) gene sequence analysis implied wide geographical circulation of a new CECoV variant. Whole genome sequencing suggested the main circulating 2022 variant was most closely related to one previously identified in 2020 with additional spike gene recombination; all variants were unrelated to CECoV-like viruses recently associated with human respiratory disease. Identifying factors that drive population-level evolution, and its implications for host protection and virulence, will be important to understand the emerging role of CECoV variants in canine and human health, and may act as a model for coronavirus population adaptation more widely.

## Introduction

Recent spillover events by coronaviruses of animal origin into humans has reminded us of the potential for coronaviruses to emerge in a population with devastating effect [1]. Subsequent evolution may create variants, partly in response to natural and, where it exists, vaccine-induced immunity. This can lead to population persistence and repeated localised outbreaks of disease, with profound implications for global health and welfare [2]. There are few natural models of disease where such coronavirus emergence and evolution can be studied.

Canine enteric coronavirus (CECoV) is an alphacoronavirus closely related to the viral cause of feline infectious peritonitis (FIPV) and transmissible gastroenteritis (TGEV) of pigs. The virus has a complex evolutionary history in which recombination has played a significant part, facilitated by the high prevalence of infection allowing for frequent mixed infections and the coronavirus genome replication strategy based on homologous recombination [3,4]. A somewhat complex nomenclature has also evolved based on immunological and more recently sequence differences in key domains of the spike gene responsible for receptor binding. Initial delineation of type I and II CECoVs was based largely on serological differences; type I CECoVs were also found to contain an additional ORF (ORF3) that is not present in either type II CECoVs or related FIPV and TGEV [5]. As more sequence data has become available, type IIb and IIc (also called type I/II) variants have been delineated based on putative recombination events in the N terminal spike domain between the original (type IIa) CECoVs and either TGEV [6,7] or type I CECoV [8] respectively [3,9]; these have been suggested to impact receptor binding and virulence although empirical evidence for this for CECoV is currently limited.

CECoV has generally been considered a cause of mild endemic gastroenteritis [10,11] with only sporadic case reports of more severe disease usually as a result of coinfection with other enteric viral pathogens [12–16]. Widespread endemic infection of a particular strain or variant has rarely been described apart from in rescue shelters [17]. However, in February 2020, an outbreak of severe vomiting and diarrhoea occurred across the UK in dogs which molecular analyses suggested may have been caused by a nationally distributed variant of CECoV [18]. In 2022, a similar outbreak of severe diarrhoea and vomiting was again being reported in the UK on social and mainstream media, both by members of the public and also veterinary professionals. Cases were initially described in coastal regions of Yorkshire; speculation about early aetiologies included contact with dead marine animals on beaches.

Here we describe an interdisciplinary response based on data science, field epidemiology, microbiology and genomics, to identify the nature and spread of this outbreak. We identify widespread distribution of what appears to be a variant of those CECoVs present in 2020 compounded by further recombination in the spike protein gene. The emergence of new CECoV variants and its implications to canine health are discussed. In addition, recent evidence of variants of CECoV in human respiratory disease [19,20], suggests a need to better understand how this observed evolution might represent a new risk to human health. We further propose CECoV can provide an accessible model to study the mechanisms and consequences of endemic coronavirus population adaptation and evolution.

## Results

### Veterinary syndromic and laboratory surveillance data

Longitudinal analysis of Main Presenting Complaint (MPC) Vet data from the UK as a whole, using a logistic latent periodic Gaussian process model adjusted for COVID-19 lockdown periods, suggested clear seasonality of canine gastroenteric disease, typically peaking in January and February each year at 4-5% of consultations (Figure 1A). In 2020, several weekly data points exceeded the 99% credible interval; it was these data points that constituted the first outbreak previously described [17]. At a national level, other years, including 2022, were within credible intervals suggestive of more normal seasonal variation. In contrast, whilst data for Yorkshire showed similar overall seasonality, two weekly data points also exceeded the 99% credible interval in 2022 (weeks commencing 10^th^ and 17^th^ January) with the following week (week commencing 24^th^ January) exceeding the 95% credible interval; this was considered to be consistent with an outbreak (Figure 1B). Diagnostic results from the laboratory data showed a similar seasonal profile for CECoV diagnoses, with the proportion of tested samples that were testing positive peaking over 0.2 each winter in January or February (Figure 1C).

**Figure 1.**
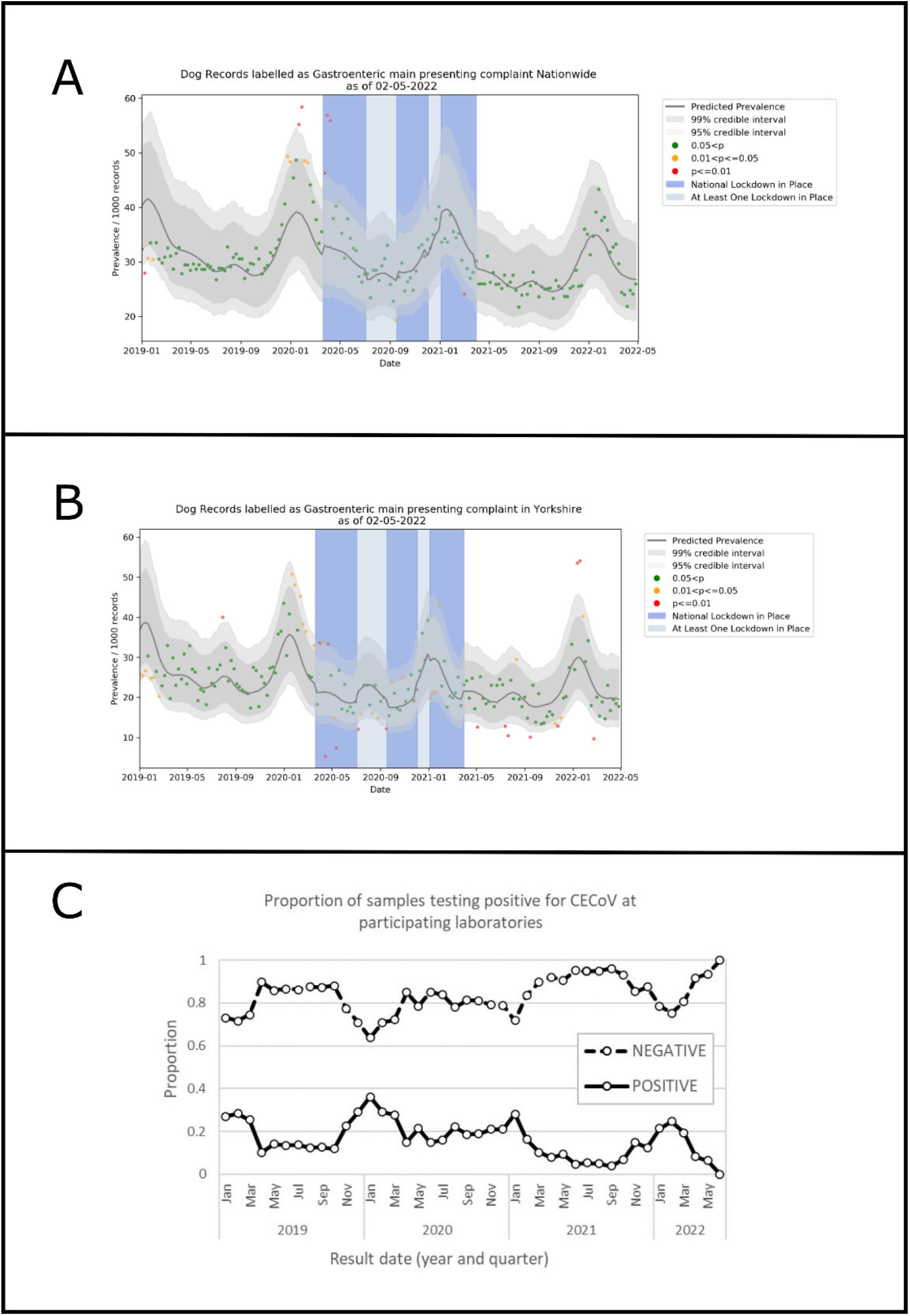
Yorkshire (A) and National (B) temporal distributions of consultations classified by the attending practitioner as mainly associated with gastroenteric disease. C) Temporal distribution of the proportion of CECoV samples testing positive and negative from diagnostic laboratories.

### Questionnaire data

In total, 28 responses were received from veterinarians caring for animals (20 cases, 8 controls), and 438 from owners of cases. The main clinical signs reported by both were vomiting, diarrhoea and inappetence. Most cases lasted between 24-48 hours (n=3, 15.0% of vet responses: n=63, 14.4% of owner responses), or 3-7 days (n=7, 35.0% of vet responses: n=170, 38.8% of owner responses). Many cases were described as still ongoing (n=9, 45% of vet responses: n=78, 17.8% of owner responses). Of the owner reported cases, 182 (41.6%) came from a multidog household; of these co-habiting dogs, 108 (59.3%) also showed gastrointestinal signs of vomiting and/or diarrhoea.

A potential link to beach walking was expressed early in the outbreak; only five (25%) of the 20 cases reported by vets said the patient had recently visited a beach; this compared to two (25%) of the eight controls. Broadly similar profiles of feeding were reported by both veterinary- and owner-reported cases and veterinary-reported controls with commercial dog food most common; feeding of raw diets was rare. None of the cases (or controls) reported contact with COVID cases in the household. The vast majority of cases were vaccinated (all 20 of the veterinary cases and 395 (90.2%) of the owner cases), and although vaccine details were not collected, this likely included protection against canine parvovirus, the main known infectious cause of severe vomiting and diarrhoea in dogs [21].

### Sample analyses

Faecal samples received at the University of Liverpool included 46 direct from veterinary practices (45 cases, 1 control). These were supplemented with 87 samples from dogs sent for a commercial diagnostic PCR panel test (IDEXX Laboratories, Wetherby, UK), 16 (18%) of which tested positive for CECoV at the laboratory.

A total of 18 of the 46 samples (39.1%) received directly from veterinarians tested positive by M (membrane) gene PCR. Of the 87 laboratory samples received from the virtual biobank, a random 27 (31.0%) were screened using the M gene PCR. Amplicons of the correct size were obtained from 19 diagnostic samples including all 16 that originally tested positive in the laboratory, and three that tested negative.

Thirty-six of these 37 amplicons were successfully Sanger sequenced producing M gene sequences of between 288 and 323 nucleotides in length (Figure 2A). These were supplemented by 19 sequences from dogs submitted to University of Lincoln in a parallel study. The University of Liverpool 2022 samples clustered as one main variant (25/36 sequences, 100% identity). The remaining 30 sequences were distributed into 14 minor variants with between 86.9% and 98.9% identity to the main 2022 variant. The 2022 main variant shared 98.9% sequence identity to the main 2020 variant, with both being widely distributed across the UK (Figure 2B).

**Figure 2.**
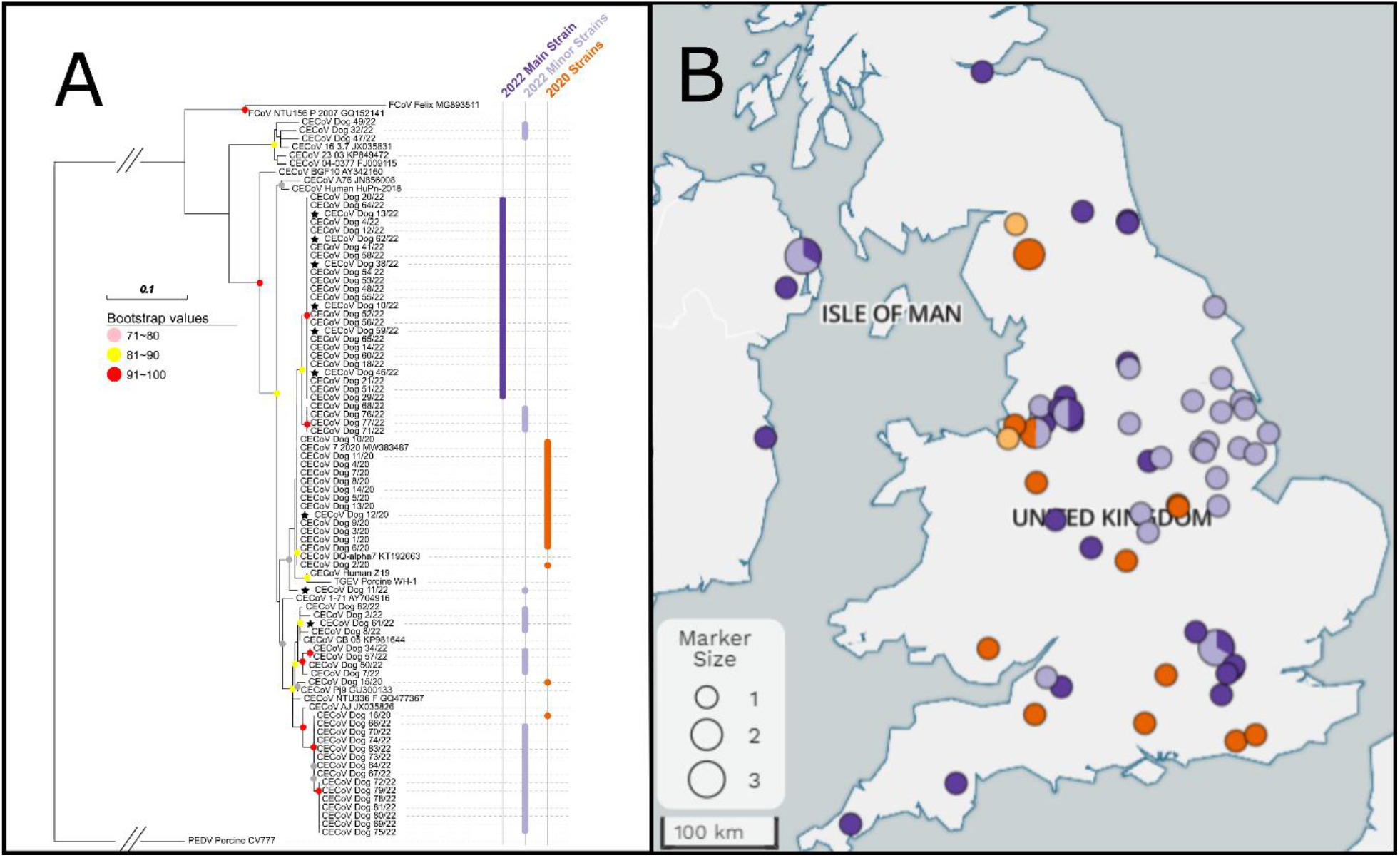
(A) Maximum-Likelihood tree of partial M gene (315bp) sequences. The scale represents the number of base differences per site. Sequences obtained from samples collected in this study (2022) are marked in purple (main strain) and blue (minor strains). Sequences obtained from 2020 are marked in orange. Samples that have also been whole genome sequenced as part of this study are indicated by stars. (B) Geographical distribution of CeCoV M variants across the UK corresponding to colours in A. Each location identifier can be linked to the phylogeny by going to an interactive interface at https://microreact.org/project/uZuCTLwtXkNQxpodQeZFS6-cecov-m-gene-2022.

### Whole genome sequencing

Initial amplicon tiling experiments based on reference genomes from previous outbreaks provided only partial genomes (between 3.5-86% breadth of coverage at >10X depth; mean 42.6%), with genome mismatches likely to be associated with failure of several primers from the initial (version 1) tiling scheme attempt. Subsequent Sequence Independent Single Primer Amplification (SISPA) allowed for the recovery of four complete genomes (Dogs 10/22, 11/22, 61/22 and 12/20). For the main circulating 2022 strain (Dog 10/22); this sequence was used as a reference to redesign a new primer scheme (version 2) resulting in the production of a further six near full length genomes.

Sequences for the major variant from 2022 (e.g. Dog 10/22) were more closely related to the major variant from 2020 (Dog 7/20 [MT906864]) over the majority of the genome (96% coverage, 97.08% identity) than to other UK viruses from either 2020 or 2022 for which near whole genome sequences were available (100% coverage, 92.66% identity; dog 15/20 [MT906864]). The lower coverage for the match between Dog 10/22 and Dog 7/20 was associated with very low sequence similarity in the S1 domain (5’ region) of the spike gene (Figure 3). This mismatched area was closely related to A76-type viruses suggestive of a recombination event which was confirmed by Gubbins (Supplementary Figure S1) at a *p* cut off of 0.05/number of substitutions occurring on the branch.

**Figure 3.**
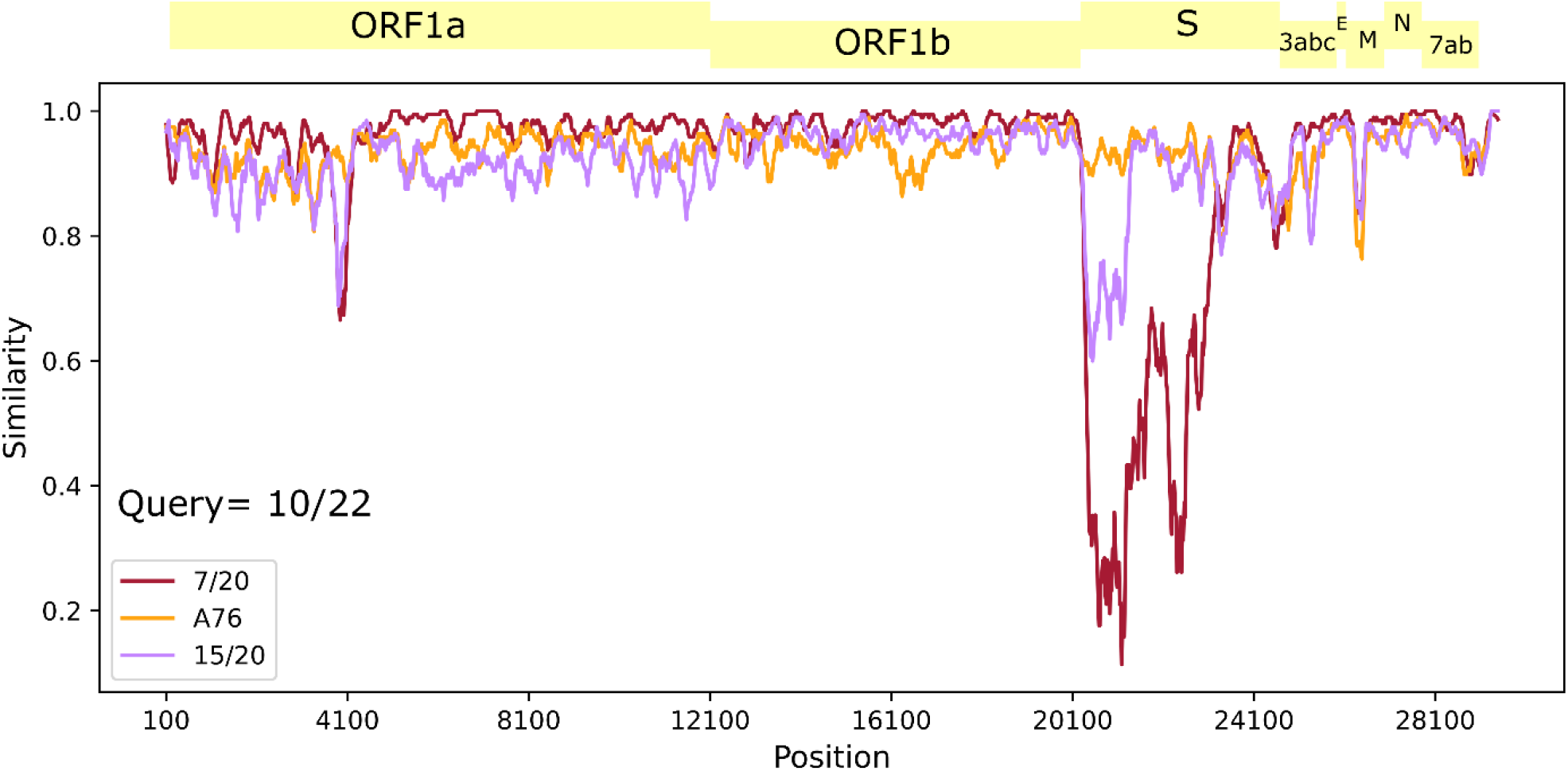
Simplot analysis using the main variant observed in the 2022 UK outbreak (Dog 10/22) as a reference compared with the main and minor strains from the 2020 outbreak (Dog 7/20 and 15/20 respectively) and the A76 strain. S=spike, E=envelope, M=membrane, N=nucleocapsid. Spike gene and core genome phylogenetic reconstruction.

Due to these perceived recombination events within the spike gene, two distinct approaches were taken to explore the evolutionary histories of CECoVs in the UK. In the first, separate phylogenetic reconstructions were undertaken for each of the spike gene regions under various selection pressures, namely the NTD and C subregions of the S1 domain (both of which have independent cell receptor binding functions), and the S2 domain (responsible for cell fusion) (Figure 4). Of particular relevance to this study, the recombinant nature of the spike protein gene from CECoV major 2022 variant from Dog 10/22 was confirmed, clustering closely with FCoV/CECoV serotype 1 strains in the S1 NTD region (Figure 4B) but with FCoV/CECoV serotype 2 strains in the S2 domain (Figure 4D). A similar recombinant serotype I/II recombinant spike protein pattern was observed for a minor variant from the 2020 outbreak (Dog 15/20).

**Figure 4.**
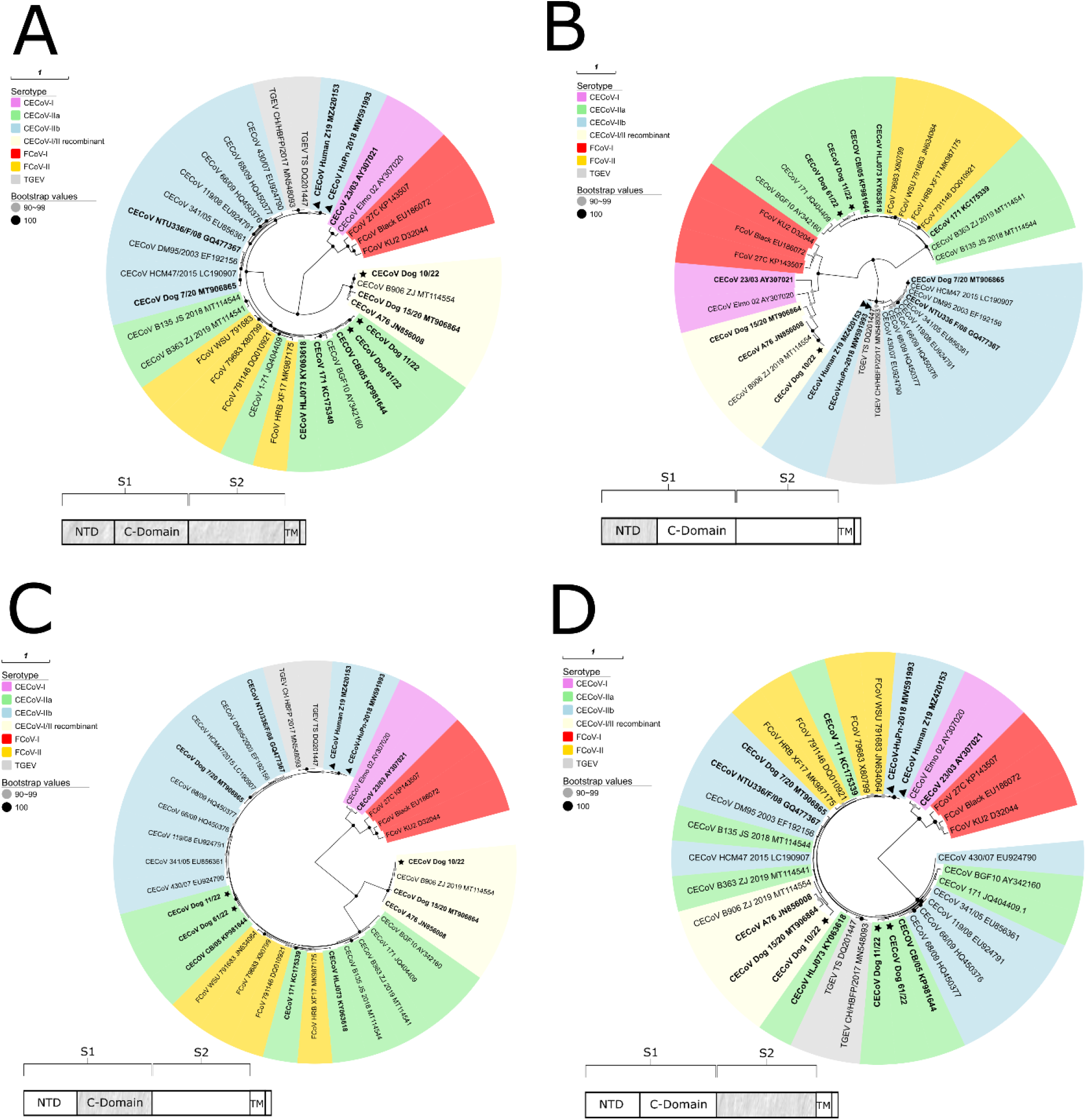
Panel of S-gene maximum-likelihood phylogenies for serotype I/II CECoV, Feline coronavirus (FCoV) and Transmissible gastroenteritis virus (TGEV) as well as recombinant strains. A) Full Spike gene. B) N-Terminal subregion (NTD) of S1 domain. C) C subregion of S1 domain. D) S2 domain. Stars indicate 2022 UK strains, triangles indicate human pneumonia strains. Bold leaves correspond to taxa used in Figure 5 core genome phylogeny. TM = transmembrane domain.

The second approach taken involved masking putative recombinant regions (Supplementary Figure S1) to construct a core genome phylogeny of 2022 CECoV UK isolates, alongside other archived strains. The main 2022 circulating variant from this study (Dogs 10, 13, 38, 46, 59 and 62) had very little within strain diversity (number of nucleotide differences per site (π) = 0.00049) and shared a recent common ancestor with the major variant from 2020 (Dog 12/20 and Dog 7/20 [MT906865]) (Figure 5). In contrast, two of the four genomes generated via the SISPA method (Dogs 11/22 and 61/11) formed a clade with the minor variant of 2020 (MT906865). Notably, the close clustering of 10/22, 15/20 and A76 seen in the spike gene (Figure 4) was not evident at the core genome level.

**Figure 5.**
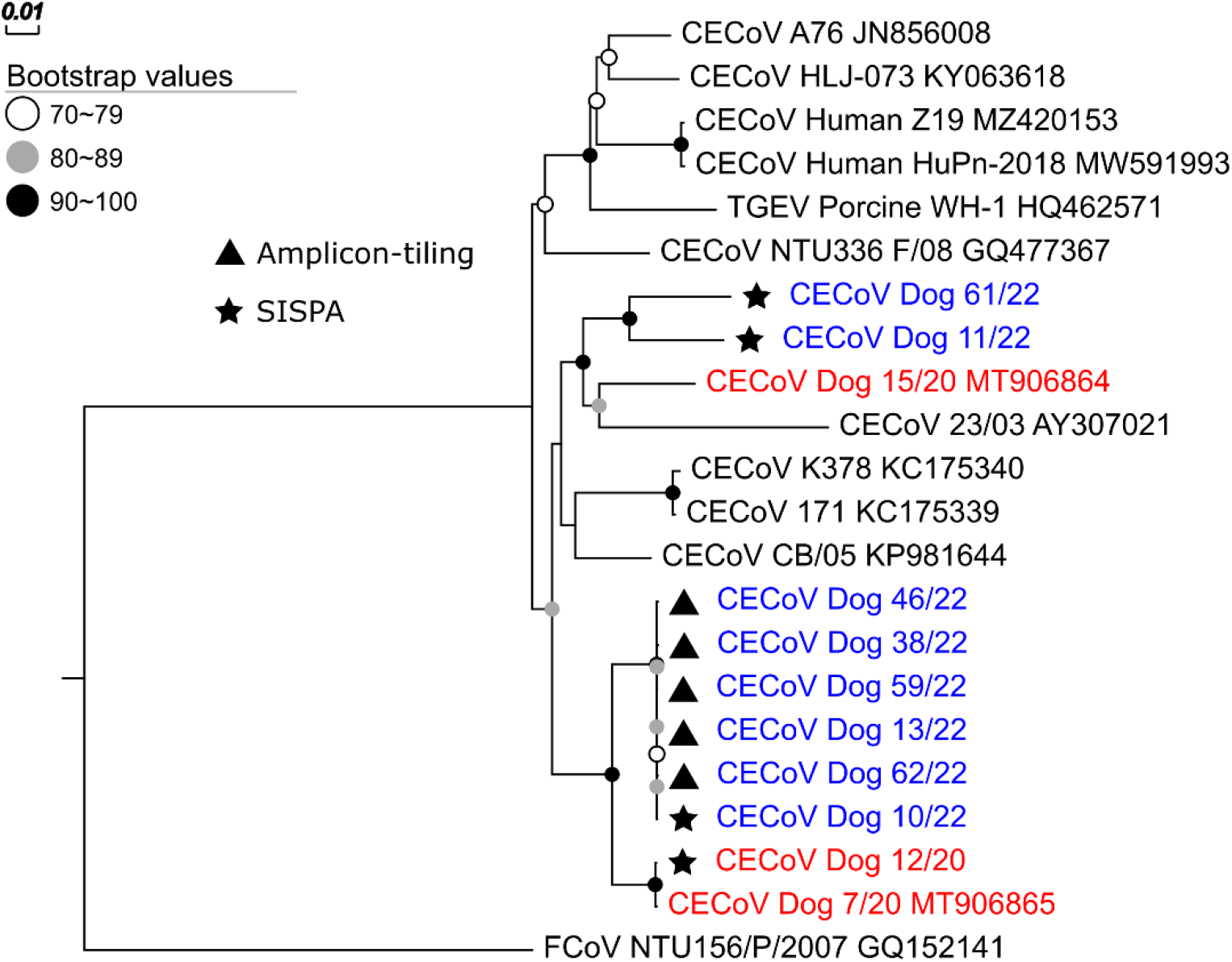
Maximum-likelihood phylogenomic analysis (final alignment 23,632 base pairs) of CECoV core genomes from 2020 (red) and 2022 (blue) UK outbreaks alongside the closest matching Genbank matches. Sequences identified in this study are indicated by stars (SISPA) and triangles (amplicon-tiling). The spike protein gene and other areas of recombination, as detected by Gubbins, were excluded from analysis.

Serotype IIb strains with evidence of causing human pneumonia (HuPn-2018 and CECoV-Z19) were not closely related to any of the circulating UK strains from this study which grouped with serotype IIa or as the I/II recombinant (Figure 5). Although the 2020 UK major strain (Dog 7/20 [MT906865]) was classed into IIb, alignment of amino acids of the spike NTD region hypothesized to lead to a tropism shift from enteric to respiratory tissues demonstrated this isolate likely lacks the amino acid changes responsible for this transition (Figure 6).

**Figure 6.**
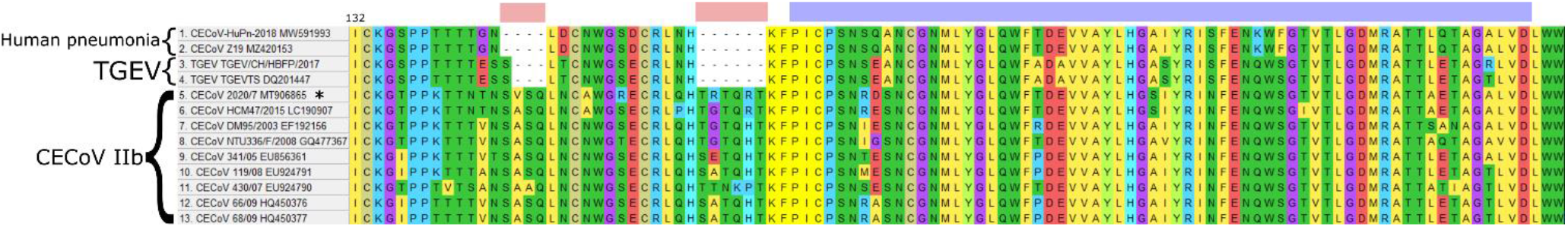
Amino acid sequence alignment of the S1 NTD region (modified from Zehr et al. [19]). The blue bar represents the area identified experimentally as being pertinent to sialic acid binding (a viral tropism determinant) in TGEV. The red bars represent upstream deletions which could also potentially impact sialic binding affinity in human pneumonia and TGEV alphacoronaviruses. Asterisk indicates the major UK 2020 variant (Dog 7/20) previously identified by the authors [18]. The number at the beginning of the alignment indicates amino acid position.

## Discussion

Coronaviruses persist and evolve over multiple annual cycles with profound implications for immunity, vaccination, disease emergence and population health. Here we took a rapid and multidisciplinary approach to a seasonal outbreak of gastrointestinal disease in dogs. Results of health record analysis clearly showed a repeated winter rise in gastrointestinal disease, statistically beyond previous seasonal credible intervals for Yorkshire. Diagnostic laboratory data suggested this may be associated again with CECoV infection. Questionnaire responses helped refute links to possible exposures to beaches or COVID cases and highlighted the severity and prolonged duration of many cases. Sequence analysis identified diverse variants of CECoV circulating in UK dogs, with one predominating. Whole genome sequencing showed this variant was closely related to that which was associated with a similar outbreak two years earlier [18] with additional spike gene recombination. This ongoing evolution is likely to be contributing to the observed seasonal pattern of canine disease and represents a valuable model system in which to study coronavirus population adaptation over repeated years.

Canine and other companion animal populations historically lack structured population health surveillance as is generally considered the norm for humans and food animal populations. We have been developing a national surveillance system based on reusing Electronic Health Records (EHRs) from animals presenting to a sentinel population of UK veterinary practices. We have shown how the addition of a practitioner derived syndrome code can add new granularity into, for example, the use of critical antibiotics [22], as well as spot important changes in the pattern of disease that enabled us to identify and respond to an outbreak of gastrointestinal disease in 2020 [18]. As apparent from the empirical estimates of Main Presenting Complaint (MPC) prevalence in Figure 1, case data is highly variable and detecting the presence of an “outbreak” in the presence of such noise is challenging. The greatest source of variation may be assumed to be due to changes in the total number of consultations occurring each day, which is largely affected by human behaviour.

This was certainly apparent during periods of COVID-19 social distancing restrictions [23], though the addition of a term into our model appears to have adequately adjusted for the otherwise artificially high empirical prevalence otherwise observed, allowing the filter distribution of the “business as usual” number of cases still to highlight weeks in which MPC cases were unusually high or low. One limitation of the present study was the inability to collect sufficient control data, such that our field epidemiology analysis was largely descriptive only. That said, such an approach did allow for the rapid evaluation of media concerns about changing patterns of disease, with our statistical validation confirming the presence of outbreaks. This highlights how such a system can be utilised for surveillance in such previously neglected populations.

As well as data from veterinary practitioners, the growing collaboration with diagnostic laboratories has allowed us to identify previously unseen patterns in pathogen diagnosis. Consistent with previous findings [18], the proportion of diagnostic samples testing positive for CECoV was shown to have a more or less consistent seasonality, peaking again in January 2022 coincident with the rise in gastroenteric disease seen with the MPC data. Such links, whilst not proof of any association, clearly can provide a rapid route to more targeted investigations such as PCR and sequencing as employed here. In this case, we were able to collect samples both directly from owners and veterinarians caring for suspect cases, as well as from a diagnostic laboratory, reusing clinical samples before they are discarded; we have termed this approach a virtual biobank [24,25] and suggest that where such collaborations can be developed and maintained, they can represent another component of surveillance particularly in resource-poor, data-rich populations like companion animals.

Two approaches were taken here to sequence analysis. The first relied on conventional PCR of the conserved coronavirus M gene based on a published cross-reactive assay [26] allowing for rapid assessment of CECoV diversity. Over the 2022 sampling period, fifteen sequence types were identified, with one predominating. Clearly the method used here can’t formally link CECoV infection to disease. However, we argue the dominance of the prevalent CECoV population by a single variant suggestive of ecological advantage, coupled with spike gene mutations likely to impact transmissibility or immunity means if we used a similar nomenclature to that used for SARS-CoV2 [27], we might classify CECoV 10/22, as a variant of interest to the dog population.

Advances in genome sequencing technologies have allowed for high-throughput, fast and cost-effective means of generating viral genomes for disease surveillance, giving more complete appreciation of virus evolution including by both substitution and recombination [2]. In the recent SARS-CoV-2 pandemic, several assays were developed based on an amplicon-tiling method using circulating variants as references to update primer schemes as and when they emerged [28]. Here, a tiling approach based on genome sequences available from 2020, gave incomplete sequences, likely due to primer mismatch. In this context, such as might occur at the beginning of an outbreak where preexisting diversity may be poorly catalogued, a SISPA approach can be more resilient to sequence mismatches as preliminary template enrichment is sequence agnostic. Once a SISPA-derived genome sequence became available, then a variant-specific tiling protocol was successfully deployed. As such, an iterative protocol (PCR screening if available; targeted SISPA; variant-specific amplicon tiling; back to SISPA where amplicon dropouts occur; Supplementary Figure S2) represents an efficient sequencing strategy for the progression of outbreaks or for use in relatively resource poor, neglected populations where prior knowledge of circulating variants will likely be limited.

Preliminary sequence similarity analysis suggested a recombination event in the recent origin of 10/22 which was subsequently confirmed by the Gubbins algorithm; the subsequent comparison of the core genome and spike protein gene phylogenies painted a complex picture. Although major variants from 2020 and 2022 UK outbreaks (Dog 7/20, Dog 10/22) shared recent common ancestors on a core genome phylogeny, the spike protein genes showed radically different evolutionary origins. For example, the minor variant for 2020 (Dog 15/20), strain A76 (JN856008) and the major variant from this study (Dog 10/22) all possessed a recombinant serotype I/II spike gene (referred to elsewhere as IIc), first reported in Sweden [29] but now seen more widely [18,30–31]. In contrast, the core genome phylogeny grouped these strains into separate clades, consistent with homologous recombination. Thus, the most parsimonious explanation for recent evolutionary events in circulating CECoV UK strains, is past co-infection of a recombinant serotype I/II strain (e.g., Dog 15/20, A76) and a virus similar to the major variant strain from 2020 (Dog 7/20), leading to a hybrid containing the serotype I/II spike gene and the core genome from Dog 7/20. The recent identification of viruses nearly identical to dog 7/20 in Raccoon dogs in Wuhan China suggests a need to include wildlife sampling in the search for the origin of such recombinants [32]. Regardless, the complex mosaic nature of CECoV genomes suggests that whole genome sequencing is required for future surveillance; a focus on the S gene, although relevant for understanding disease manifestation, cannot by itself be used to determine historic spread of CECoVs.

Due to the role of the spike protein in determining receptor binding, host affinity, immunoevasion and severity of disease [3], it is expected these recombinant variants will behave differently to prototypical CECoV strains. Indeed, evidence from other coronaviruses suggest a remarkable plasticity, and even a redundancy, in receptor binding domains, thereby facilitating new receptor acquisition [33]. Specifically, through *in-vitro* work, Regan et al [8] demonstrated the recombinant serotype I/II strain A76 possesses an altered and highly canine-specific receptor-binding profile, able to infect multiple dog-cell lines in contrast to type II viruses that could also infect feline cells. In TGEV, mutations in the NTD region of the spike gene associated with loss of sialic acid co-receptor interactions are linked to loss of enteric tropism [34].

Betacoronaviruses (e.g., SARS-CoV-1, SARS-CoV-2 and MERS-CoV) have been the subject of increasing scrutiny due to several spillover events leading to human respiratory infections. Such spillover events are however not limited to the betacoronaviruses with structured sampling of bat coronaviruses identifying probable ancestors of human alphacoronaviruses [35]. Of great relevance here is the recent identification of a CECoV-like virus (serotype IIb) in human pneumonia cases from Malaysia [36] and independently from the urine of someone returning from Haiti [37], suggesting this phenomenon may be more commonly occurring. The precise mechanism for such spillover events is not known but it is reasonable to speculate this will include spike gene mutations; indeed, replacement of the spike gene of the betacoronavirus mouse hepatitis virus (MHV) with that of feline coronavirus (FCoV), allowed the MHV recombinant to acquire permissiveness to cats [38]. Sequence analysis by Zehr et al [19] identified multiple recombination events between CECoV and FCoV; temporal analysis suggested CCoV-HuPn-2018 may have diverged from a lineage most recently circulating in cats between 1846 and 1976 (95% HPD—Highest Posterior Density Interval), with a median estimate of 1957 [19]. The lack of similarity between 10/22 and CECoV-HuPn-2018 and lack of conserved deletions/unique substitutions between 7/20 and CECoV-HuPn-2018 putatively attributed to loosened sialic acid binding, suggests there is currently unlikely to be a zoonotic threat from these highly prevalent CECoV variants of interest. That said, close interaction between dogs, cats and owners, the rapid evolution of these viruses, coupled with an ongoing paucity of data for CECoV diversity in dogs (and cats), argues for a pressing and ongoing need to monitor infection with these viruses both in pets and humans.

In conclusion, detailed surveillance has allowed us to describe a profound seasonality to CECoV associated with sporadic larger outbreaks. In-depth sequence analysis shows outbreaks are coincidental with the emergence of widely distributed CECoV variants with diverse spike genes, including recombinant I/II serotypes, which are likely to be associated with varying virulence, transmissibility, host-specificity and/or spillover potential. Further studies are urgently needed to assess the virulence of the identified variants of interest and provide greater depth of sequence information for these canine (and feline) populations and the coronaviruses they harbour. Recent identification of CECoV variants in Raccoon dogs from Wuhan closely related to the major variant we identified in 2020 [32] heightens the need for efficient and continued surveillance of circulating variants in pet dogs, wildlife and humans, enabled by efficient high-throughput genomic techniques, to understand the respective roles these strains play in both animal and human health.

## Methods

### Veterinary syndromic and laboratory surveillance data collection

Electronic health data from January 2019 to present were obtained from SAVSNET (the Small Animal Veterinary Surveillance Network) from sentinel networks of participating veterinary practices (Vet data) and diagnostic laboratories (Lab data). Vet data is collected in real time from approximately 10% of UK veterinary practices and for this study included consultation date, owner postcode mapped to NUTS2 levels of spatial resolution, species of animal, as well as a practitioner-derived classification of the main presenting complaint (MPC) [39]; for the MPC, the attending practitioner chooses one of 10 syndromes that describes the main reason the animal has been bought to the surgery (e.g., gastroenteric disease, respiratory diseases, pruritus, trauma, tumour); in the context of this work, we focus on those where the chosen MPC was gastroenteric disease.

For the Laboratory data, SAVSNET receives test results directly from the diagnostic laboratories that performed them; approximately 60% of the veterinary practices in the UK submit samples to these participating laboratories. Received data includes the pathogen being tested for, the assay methodology, result and date (variably sample submission or reporting date), and the full postcode of the submitting veterinary practice [25].

### Temporal analysis of Major Presenting Complaint (MPC) data

*y*_*t*_ MPC cases out of a total of *N*_*t*_ consults in week *t*=1,…,174 were collected over a 41 month period spanning 1st January 2019 to 2nd May 2022. We modelled *y*_*t*_ as a Binomial random variable such that

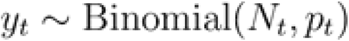

where *p*_*t*_ is the probability of a consult in week *t* being an MPC, with the log odds of being an MPC in week *t* modelled as a linear combination of terms as described below.

During our time period of interest, the presence of social distancing (lockdown) restrictions due to COVID-19 had a marked effect on the apparent prevalence of MPCs. The cancellation of routine consults (e.g. vaccinations and health checks) reduced *N*_*t*_ for weeks during which social distancing restrictions applied, though emergency consults for gastroenteric disease appear to still have taken place [23]. The result of this was to increase the apparent prevalence of MPC during the affected weeks. To capture this effect, we introduced a dummy variable, *z*_*t*_ taking the value 1 if week *t* was affected by social distancing and 0 otherwise [40].

We then let

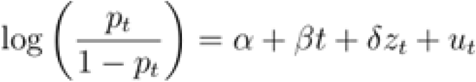

where *α* is the mean log odds of an MPC consult, *β* represents a linear time trend capturing long-term drift in MPC prevalence (the effect on the log odds of MPC for a 1 week increase in time), *δ* represents an offset in the log odds for MPC for weeks in which social distancing was imposed, and *u*_*t*_ represents a time-varying random effect.

The random effect *u*_*t*_ allows us to model periodic serial correlation in our weekly observations, as well as any extra-Binomial variation that might contribute to the overall variability of cases from one week to the next. We model the vector ***u*** as a Gaussian process with mean 0 and covariance matrix *Σ*^2^ such that

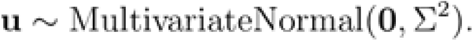

The covariance matrix *Σ*^2^ captures the correlation between two variates *u*_*t*_ and *u*_*s*_ spaced *s-t* weeks apart, and we assume the correlation follows a periodic function

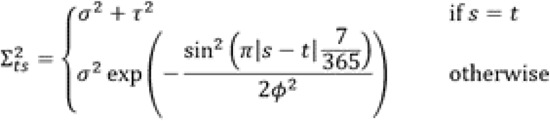

Here, *σ*^2^ represents the variance between two timepoints spaced a year apart, *ϕ* represents the lengthscale of the correlation (essentially how correlated any two adjacent timepoints are), and *τ*^2^ represents extra-Binomial variability due to observation error. For identifiability reasons, we fix *ϕ*=0.32year^−1^, tuned manually to give a satisfactory amount of smoothing over the timeseries. The model was cast in a Bayesian setting, for which prior distributions were used for all unknown parameters *α, β, δ, σ, τ* (Table 1).

**Table 1:** Prior distributions used for the longitudinal latent Gaussian process model

| Parameter | Prior Distribution |
| --- | --- |
| $\alpha$ | Normal(0,1000) |
| $\beta$ | Normal(0,100) |
| $\sigma^2$ | HalfNormal(5) |
| $\tau^2$ | HalfNormal(5) |
| $\delta$ | Normal(0,100) |

The model was fitted using the No-U Turn Sampler implementation in the Python package PyMC3 for 6000 iterations with a 1000 iteration burn in period, to provide numerical estimates of the joint posterior distribution [41].

The advantage of Bayesian inference in our context is that it allows us to compare our observations *y*_*t*_, *t=1*,…,*174*, with the filter distribution

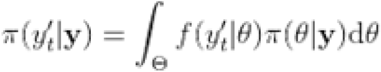

For each observation, we calculate

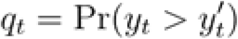

highlighting timepoints where 0.95<*q*_*t*_*<*0.99 as possibly higher than expected, and where *q*_*t*_*>0*.*99* as likely to be higher than expected. Conversely, we identify cases where 0.025 < *q*_*t*_ < 0.05 and *q*_*t*_ < 0.025 as possibly or likely to be lower than expected respectively.

### Field epidemiology

To rapidly collect more detailed descriptive data on potential cases of vomiting and diarrhoea, a questionnaire was developed and trialed for circulation to owners and veterinary surgeons. This validated questionnaire contained information on the owner’s location, the type of dog, its vaccination status, the nature of any disease including its onset, places the dog was exercised, feeding habits, and contact with other animals, including people in the household with COVID-19. Equivalent data was also requested for control animals. Owners and veterinarians were recruited by convenience both by the SAVSNET website, and also by social media releases in both the veterinary and public domains. These data were analysed descriptively.

### Dog faecal sample collection

Samples for microbiological analysis were obtained from two sources. Firstly, veterinary surgeons recruited via the SAVSNET website (https://www.liverpool.ac.uk/savsnet/dog-gi-investigation/) were asked to submit a sample of voided faeces to the University of Liverpool from dogs presented to them with either vomiting and/or diarrhoea of unknown aetiology or from control dogs with no such symptoms. Secondly, canine faecal samples sent directly to participating diagnostic laboratories to be tested for a panel of canine enteric pathogens were also retrieved from the SAVSNET “virtual biobank” [23]. All data and sample collection was approved by the University of Liverpool Research Ethics Committee (Virtual Biobank samples, RETH00964) or the Veterinary Research Ethics Committee (Veterinary samples, VREC922ab).

### PCR and sequencing

To extract total nucleic acid (TNA) for downstream molecular analyses, approximately 30 mg of faeces was suspended in PBS at a ratio of 10% (w/v) and homogenised for 30 seconds. The mixture was centrifuged at 13,000 rpm for 5 mins and 140 µl of supernatant used for RNA extraction (Qiagen viral RNA kit). The final TNA was eluted in 60 µl of molecular grade water and stored at −80°C.

For rapid screening of samples of CECoV by PCR and Sanger sequencing, extracted RNA was reverse transcribed using random hexamers and Superscript III (Thermo Fisher Scientific). Amplification of the partial CECoV M gene (409 nt) was attempted using primers CCV1 and CCV2 with minor modification [25]; the PCR was run as a single-stage PCR rather than as the published nested reaction to reduce the risk of false positives. All resultant amplicons were purified (Qiagen PCR purification kit) and sequenced bidirectionally using the PCR primers (Source Bioscience). Forward and reverse sequences were aligned and primer regions trimmed using ChromasPro (Technelysium: http://technelysium.com.au/wp/chromaspro/).

In order to compare different approaches to generating whole genome sequences rapidly in the face of an outbreak that was initially of uncertain cause, two different approaches to library preparation were tried. Sequence-Independent Single Primer Amplification (SISPA) is an approach that seeks to enrich for viral sequences by random amplification of short sequences of cDNA. SISPA was performed as described previously Greninger et al [42] with a few modifications. Briefly, 5μL TNA was incubated at 65°C for 5 minutes with 40pmol primer Sol-A (5’— GTT TCC CAC TGG AGG ATA NNN NN—3’) and 1 μL 10mM dNTP Mix before reverse transcription using SuperScript IV Reverse Transcriptase (Thermo Fisher Scientific) following manufacturer’s instructions with the addition of the ribonuclease inhibitor RNaseOUT (Life Technologies). Sequenase Version 2.0 (Thermo Fisher Scientific) was then utilized for second strand synthesis. cDNA was then cleaned using AMPure XP beads in a 1.8:1 ratio and eluted in 40μL nuclease-free water. Random amplification was performed on each sample using Q5 High-Fidelity 2X Master Mix (New England Biosciences) and 100pmol primer Sol-B (5’—GTT TCC CAC TGG AGG ATA —3’). A PCR assay was then undertaken using the following conditions: initial denaturation for 30 seconds at 98°C, followed by 35 cycles of 98°C for 10 seconds, 54°C for 30 seconds, and 72°C for 1 minute, with a final extension step of 72°C for 10 minutes. Amplicons were then purified using AMPure XP beads in a 1.8:1 ratio and eluted in 40μL nuclease-free water. Amplified cDNA was quantified using a Qubit 1X dsDNA HS hit and Qubit Flex Fluorometer (Thermo Fisher Scientific) with fragment lengths assessed using the Agilent 2100 Bioanalyzer system and High Sensitivity DNA Kit (Agilent). Library preparation was performed as previously described by Gauthier et al. [43]. Five samples were multiplexed on each MinION flow cell, with the addition of a negative control (Nuclease-free water) to each library. Samples were sequenced on FLO-MIN106D flow cells (R9.4.1 chemistry) on a GridION sequencing device for 72 hours using MinKNOW (Version 3.6.5) with live base calling disabled.

In contrast to SISPA, amplicon-tiling seeks to specifically enrich, by overlapping PCRs, genomes of interest in clinical samples [28]. When successful, such an approach can be cheaper and more efficient route of target genome recovery. An initial attempt (version 1) was made to use an amplicon-tiling primers based on reference sequences for CECoV genomes from the 2020 UK outbreak (MT906864 and MT906865) using PrimalScheme v1.3.2 [28]. Unfortunately, amplicon dropouts meant this method was only able to generate partial genomes. As a result, a version 2 primer scheme was based on the genome recovered from Dog 10/22 via the SISPA method which, according to the M gene phylogeny from initial screening, was the predominant 2022 variant. PCR conditions for the version 2 primer schemes were as described by Quick et al. [28]. For details of primer schemes, reference genomes and version 1 (pilot) data, see https://github.com/edwardcunningham-oakes/CECoV-outbreak-2022. Library preparation of targeted PCR amplicons was adapted from the ncov-2019 sequencing v3 (ARTIC) protocol by Josh Quick (https://www.protocols.io/view/ncov-2019-sequencing-protocol-v3-locost-bh42j8ye) with slight modifications as follows: PCR product (amplicon size 1200bp) was normalized to 100ng per sample for end-preparation and adaptor ligation steps. The ONT Native Barcoding Ligation kit (EXP-NBD196) was used for multiplexing no more than 30 samples per flow cell. All samples were sequenced on FLO-MIN106D flow cells (R9.4.1 chemistry) on a GridION sequencing device for 72 hours using MinKNOW (Version 3.6.5) with live basecalling disabled.

### Bioinformatics

Basecalling of SISPA and amplicon-tiling Fast5 files was undertaken using Guppy v4.2.2. Outputted FASTQ files were demultiplexed using PoreChop v0.2.4 [44] and quality filtered/primer trimmed using Nanofilt version 2.8.0 (size selection: 150-1500bp (SISPA), 1000-1400bp (amplicon-tiling); Average Q score: ≥ 15; Head and tail trimming: 18 bases (SISPA), 27 bases (amplicon-tiling)) [45]. Filtered and trimmed SISPA FASTQ files were uploaded to the online BugSeq v1 portal (https://bugseq.com/ upload date: Mar 29 2022) for metagenomic classification using the RefSeq database [46] with classification results summarised and viewed in Recentrifuge [47]. Reads classified as alphacoronavirus were extracted using seqtk version 1.3-r106 (https://github.com/lh3/seqtk). BLASTn was then used on a subset of the SISPA and amplicon-tiling reads to find the nearest Genbank hit to use as a reference for mapping reads to build a consensus. The mpileup option in Samtools version 1.15 [48] was used to build a consensus with areas of less than 5X coverage masked. For one genome (Dog 61/22) which had several low coverage areas (<5X depth), amplicons from the V1 amplicon tiling scheme were incorporated to generate a consensus sequence. If reference genomes were not sufficient for mapping the divergent Spike (S) gene, a custom BLAST database of canine coronaviruses S genes was used to map reads to get a consensus to be combined with the draft genome.

### Phylogenetics

M gene sequences generated by PCR and Sanger sequencing were aligned together with 2020 sequences (MT877072, MT906864 and MT906865) and others from GenBank, using ClustalW v2.1, producing an 315bp alignment in the final dataset. A Maximum-Likelihood tree was inferred in IQTree v2.2.0 [49], using default parmeters and the TIM2e+G4 model as determined by ModelFinder [50], with bootstrapping (1000 ultrafast) to give some estimate of node reliability. The M gene phylogeny and metadata were then visualised using Microreact [51] and Evolview [52].

For whole genome analysis, an initial alignment to assess for potential recombination was produced by aligning three genomes generated in this study and our previous work [18] (15/20, 7/20 and 10/22) and a reference genome (A76 [JN856008]) in MAFFT v7.490 [53]. This alignment was then visualised in SimPlot++ v1.3 [54]. All near-complete draft CECoV genomes generated using the MinION platform and their nearest Genbank matches were then aligned using the LINSI algorithm in MAFFT. Genomes were then assessed for recombination using Gubbins v2.3.4 [55], and visualised using Phandango v1.3.0 [56]. Regions with evidence of recombination were then masked using Gubbins whilst alignment regions with excessive gaps were removed using Gblocks v0.91b [57] under default parameters. This masked alignment was then used for phylogenetic analysis. For phylogenomic construction, the GTR+F+I+G4 model was used as determined by ModelFinder based on default “auto” parameters using the Bayesian information criteria [50]. A maximum likelihood (ML) tree was then estimated with IQTree v2.2.0 [49]. Finally, a phylogram was drawn and annotated using EvolView v3 [52].

Due to historic [3,32] and preliminary evidence of recombination in the spike protein (S) genes of genomes, phylogenies of the S1 and S2 domains of S genes generated in this study and other alphacoronaviruses (FcoV / CECoV I and II and TGEV) were generated using the above methods with slight modifications as follows: Pal2NAL [58] was used to ensure accurate codon alignments and the GTR+F+G4 model was used for tree construction.

## Data Availability

Raw reads, annotated genomes and M gene sequences can be found under bioproject PRJEB55544 in the European Nucleotide Archive. Primer schemes, reference genomes, pilot amplicon-tiling and BugSeq results can be found at https://github.com/edwardcunningham-oakes/CECoV-outbreak-2022.

## Acknowledgements

This work was largely funded by Dogs Trust as part of SAVSNET Agile, a project designed to develop a methodology for efficient national canine health surveillance based on health- and bio-informatics.

## Supplementary Figure Descriptions

**Supplementary Figure S1**. Phandango visualization of predicted recombination based on genome alignment of CECoV sequences. A) Functional annotation of reference genome A76, where yellow represent the full length of sequence, whilst light blue represents predicted coding sequences. B) Phylogenetic tree generated using Gubbins on recombination sites free alignment. C) Predicted regions of recombination, where red blocks represent ancestral recombination whilst blue blocks represent variant-specific recombination. D) Graph measuring SNP density.

**Supplementary Figure S2**. Proposed methodological pipeline for genome-based surveillance of canine enteric coronavirus during population outbreaks.

## References

[1] Masood, N., Malik, S.S., Raja, M.N., Mubarik, S., Yu, C. Unravelling the epidemiology, geographical distribution, and genomic evolution of potentially lethal coronaviruses (SARS, MERS, and SARS CoV-2). Front. Cell Infect. Microbiol. 10, 499 (2020).

[2] Vöhringer, H.S. et al. Genomic reconstruction of the SARS-CoV-2 epidemic in England. Nature 600(7889), 506–511 (2021).

[3] Licitra, B.N., Duhamel, G.E., Whittaker, G.R. Canine enteric coronaviruses: Emerging viral pathogens with distinct recombinant spike proteins. Viruses 6, 3363–3376 (2014).

[4] Costa E.M., de Castro T.X., Bottino F.de O., Garcia Rde C. Molecular characterization of canine coronavirus strains circulating in Brazil. Vet. Microbiol. 168, 8–15 (2014).

[5] Lorusso, A. et al. Gain, preservation, and loss of a group 1a coronavirus accessory glycoprotein. J. Virol. 82, 10312–7 (2008).

[6] Decaro, N. et al. Recombinant canine coronaviruses related to transmissible gastroenteritis virus of swine are circulating in dogs. J. Virol. 83, 1532–1537 (2009).

[7] Wesley, R.D. The S gene of canine coronavirus, strain UCD-1, is more closely related to the S gene of transmissible gastroenteritis virus than to that of feline infectious peritonitis virus. Virus Res. 61, 145–152 (1999).

[8] Regan, A.D. et al. Characterization of a recombinant canine coronavirus with a distinct receptor-binding (S1) domain. Virology, 430, 90–99 (2012).

[9] Decaro, N. et al. European surveillance for pantropic canine coronavirus. J. Clin. Microbiol. 51, 83–88 (2012).

[10] Tennant, B.J., Gaskell, R.M., Kelly, D.F., Carter, S.J. and Gaskell, C.J. Canine coronavirus infection in the dog following oronasal inoculation. Res. Vet. Sci. 51, 11–18 (1991).

[11] Keenan, K.P., Jervis, H.R., Marchwicki, R.H., and Binn, L.N. Intestinal infection of neonatal dogs with canine coronavirus 1-71: studies by virologic, histologic, histochemical, and immunofluorescent techniques. Am. J. Vet. Res. 51 (3), 247–56 (1976).

[12] Naylor, M.J. et al. Identification of canine coronavirus strains from feces by S gene nested PCR and molecular characterization of a new Australian isolate. J. Clin. Microbiol. 39, 1036–1041 (2001).

[13] Buonavoglia, C. et al. Canine coronavirus highly pathogenic for dogs. Emerg. Infect. Dis. 12, 492–494 (2006).

[14] Evermann, J.F., Abbott, J.R. and Han, S. Canine coronavirus-associated puppy mortality without evidence of concurrent canine parvovirus infection. J. Vet. Diag. Invest. 17, 610–614 (2005).

[15] Escutenaire, S. et al. Characterization of divergent and atypical canine coronaviruses from Sweden. Arch. Virol. 152, 1507–1514 (2007).

[16] Licitra, B.N., Whittaker, G.R., Dubovi, E.J. and Duhamel, G.E. Genotypic characterization of canine coronaviruses associated with fatal canine neonatal enteritis in the United States. J. Clin. Microbiol. 52 (12), 4230–4238 (2014).

[17] Stavisky J. et al. Cross sectional and longitudinal surveys of canine enteric coronavirus infection in kenneled dogs: a molecular marker for biosecurity. Infect Genet Evol. 12(7), 1419–26 (2012).

[18] Radford, A.D. et al. Outbreak of severe vomiting in dogs associated with a canine enteric coronavirus, United Kingdom. Emerg. Infect. Dis. 27, 517–528 (2021).

[19] Zehr, J.D. et al. Recent zoonotic spillover and tropism shift of a canine coronavirus is associated with relaxed selection and putative loss of function in NTD subdomain of spike protein. Viruses 14, 853 (2022).

[20] Anastasia, N. et al. Animal alphacoronaviruses found in human patients with acute respiratory illness in different countries. Emerg. Microbes. Infect. 11, 699–702 (2022).

[21] Clegg, S. et al. Molecular epidemiology and phylogeny reveal complex spatial dynamics in areas where canine parvovirus is endemic. J Virol. 85, 7892–9 (2011).

[22] Singleton, D.A., et al. Pharmaceutical prescription in canine acute diarrhoea: A longitudinal electronic health record analysis of first opinion veterinary practices. Front. Vet. Sci. 6, 218 (2019).

[23] Littlehales, R., Noble, P.M., Singleton, D.A., Pinchbeck, G.L., Radford, A.D. Impact of Covid-19 on veterinary care. Vet. Rec. 186(19), 650–651 (2020).

[24] Smith, SL, et al. A virtual biobank for companion animals: A parvovirus pilot study. Vet. Rec. 189, 556 (2021).

[25] Singleton, D.A. et al. Temporal, spatial, and genomic analyses of Enterobacteriaceae clinical antimicrobial resistance in companion animals reveals phenotypes and genotypes of One Health concern. Front. Microbiol. 12, 700698 (2021).

[26] Pratelli, A. et al. Development of a nested PCR assay for the detection of canine coronavirus. J. Virol. Methods 80 (1), 11–5 (1999).

[27] WHO. https://www.who.int/activities/tracking-SARS-CoV-2-variants. Accessed August 2022

[28] Quick, J., et al. Multiplex PCR method for MinION and Illumina sequencing of Zika and other virus genomes directly from clinical samples. Nature Protocols 12(6), 1261–1276. (2017).

[29] Escutenaire, S., et al. Characterization of divergent and atypical canine coronaviruses from Sweden. Arch. Virol. 152(8), 1507–14 (2007).

[30] Regan, A.D., et al. Characterization of a recombinant canine coronavirus with a distinct receptor-binding (S1) domain. Virology 430(2), 90–99 (2012).

[31] He, H.-J., et al. Etiology and genetic evolution of canine coronavirus circulating in five provinces of China, during 2018–2019. Microbial Pathogenesis 145, 104209 (2020).

[32] Wang, W. et al. Coronaviruses in wild animals sampled in and around Wuhan at the beginning of COVID-19 emergence. Virus Evol. 8(1), veac046 (2022).

[33] Hulswit, R.J.G., de Haan, C.A.M., Bosch, B.-J. Coronavirus Spike Protein and Tropism Changes Advances in Virus Res. 96, 29–57 (2016).

[34] Ballesteros, M.L., Sánchez, C.M., Enjuanes, L. Two amino acid changes at the N-terminus of transmissible gastroenteritis coronavirus spike protein result in the loss of enteric tropism. Virology 227(2), 378–88 (1997).

[35] Tao, T. Surveillance of bat coronaviruses in Kenya identifies relatives of human coronaviruses NL63 and 229E and their recombination history. J. Virol. 91(5), e01953–16 (2017).

[36] Vlasova, A.N. Novel canine coronavirus isolated from a hospitalized patient with pneumonia in East Malaysia. Clin. Infect. Dis. 74(3), 446–454 (2022).

[37] Lednicky, J.A. Isolation of a novel recombinant canine coronavirus from a visitor to Haiti: Further evidence of transmission of coronaviruses of zoonotic origin to humans. Clin. Infect. Dis. ciab924, (2021).

[38] Kuo L. et al. Retargeting of coronavirus by substitution of the spike glycoprotein ectodomain: crossing the host cell species barrier. J. Virol. 74(3), 1393–1406 (2000).

[39] Sánchez-Vizcaíno, F. et al. Demographics of dogs, cats, and rabbits attending veterinary practices in Great Britain as recorded in their electronic health records. BMC Vet. Res. 13, 218 (2017).

[40] The Institute for Government, 2022. Timeline of UK government coronavirus lockdowns and restrictions. https://www.instituteforgovernment.org.uk/charts/uk-government-coronavirus-lockdowns. Accessed 28 August 2022.

[41] Salvatier, J., Wiecki, T.V., Fonnesbeck, C. Probabilistic programming in Python using PyMC3. PeerJ Comp. Sci. 2, e55 (2016).

[42] Greninger, A.L. et al. Rapid metagenomic identification of viral pathogens in clinical samples by real-time nanopore sequencing analysis. Genome Med. 7(1), 99 (2015).

[43] Gauthier, N.P.G. et al. Nanopore metagenomic sequencing for detection and characterization of SARS-CoV-2 in clinical samples. PLOS ONE 16(11), e0259712. (2021).

[44] Wick, R.R., Judd, L.M., Gorrie, C.L., & Holt, K.E. Completing bacterial genome assemblies with multiplex MinION sequencing. Microbial Genomics 3(10), e000132 (2017).

[45] de Coster, W., D’Hert, S., Schultz, D. T., Cruts, M., & van Broeckhoven, C. NanoPack: visualizing and processing long-read sequencing data. Bioinformatics 34(15), 2666–2669 (2018).

[46] Fan, J., Huang, S., & Chorlton, S.D. BugSeq: a highly accurate cloud platform for long-read metagenomic analyses. BMC Bioinformatics 22(1), 160 (2021).

[47] Martí, J.M. Recentrifuge: Robust comparative analysis and contamination removal for metagenomics. PLOS Computational Biol. 15(4), e1006967. (2019).

[48] Li, H., et al. The Sequence Alignment/Map format and SAMtools. Bioinformatics 25(16), 2078– 2079 (2009).

[49] Minh, B.Q. et al. IQ-TREE 2: New models and efficient methods for phylogenetic inference in the genomic era. Mol. Biol. Evol. 37(5), 1530–1534 (2020).

[50] Kalyaanamoorthy, S., Minh, B.Q., Wong, T.K.F., von Haeseler, A., & Jermiin, L.S. ModelFinder: fast model selection for accurate phylogenetic estimates. Nature Methods 14(6), 587–589 (2017).

[51] Argimón, S., Abudahab, K., Goater R.J.E., et al. Microreact: visualizing and sharing data for genomic epidemiology and phylogeography. Microb. Genom. 2(11):e000093 (2016).

[52] Subramanian, B., Gao, S., Lercher, M.J., Hu, S., & Chen, W.-H. Evolview v3: a webserver for visualization, annotation, and management of phylogenetic trees. Nuc. Acid. Res. 47(W1), W270– W275 (2019).

[53] Katoh, K., & Standley, D. M. MAFFT Multiple sequence alignment software Version 7: Improvements in performance and usability. Mol. Biol. Evol. 30(4), 772–780 (2013).

[54] Samson, S., Lord, É., & Makarenkov, V. SimPlot++: a Python application for representing sequence similarity and detecting recombination. Bioinformatics, 38(11), 3118–3120 (2022).

[55] Croucher, N.J. et al. Rapid phylogenetic analysis of large samples of recombinant bacterial whole genome sequences using Gubbins. Nuc. Acid. Res. 43(3), e15 (2015).

[56] Hadfield, J. Phandango: an interactive viewer for bacterial population genomics. Bioinformatics 34(2), 292–293 (2018).

[57] Castresana, J. Selection of conserved blocks from multiple alignments for their use in phylogenetic analysis. Mol. Biol. Evol. 17(4), 540–552 (2000).

[58] Suyama, M., Torrents, D., & Bork, P. PAL2NAL: robust conversion of protein sequence alignments into the corresponding codon alignments. Nuc. Acid. Res. 34, W609–W612 (2006).

