## Supplementary figures and images for "Emerging variants of canine enteric coronavirus associated with seasonal outbreaks of severe canine gastroenteric disease"

### Supplementary Figure S1 2 10b.png

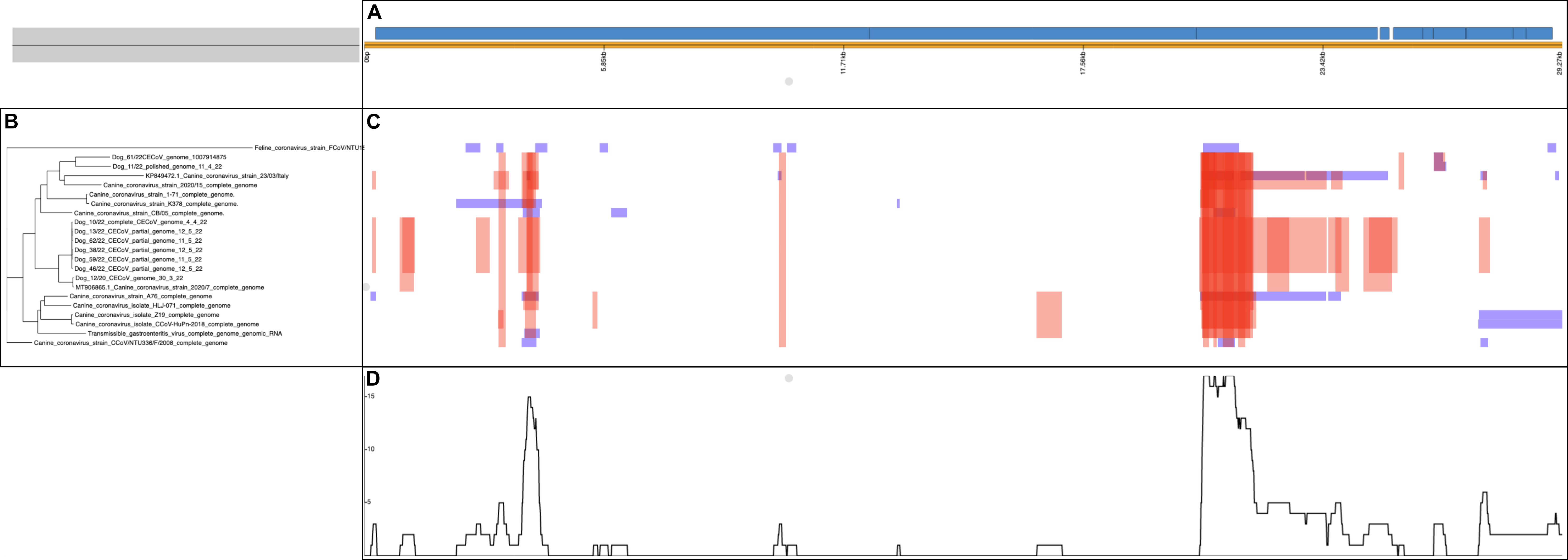

### Workflow_FigureS2_29_9_22b.png

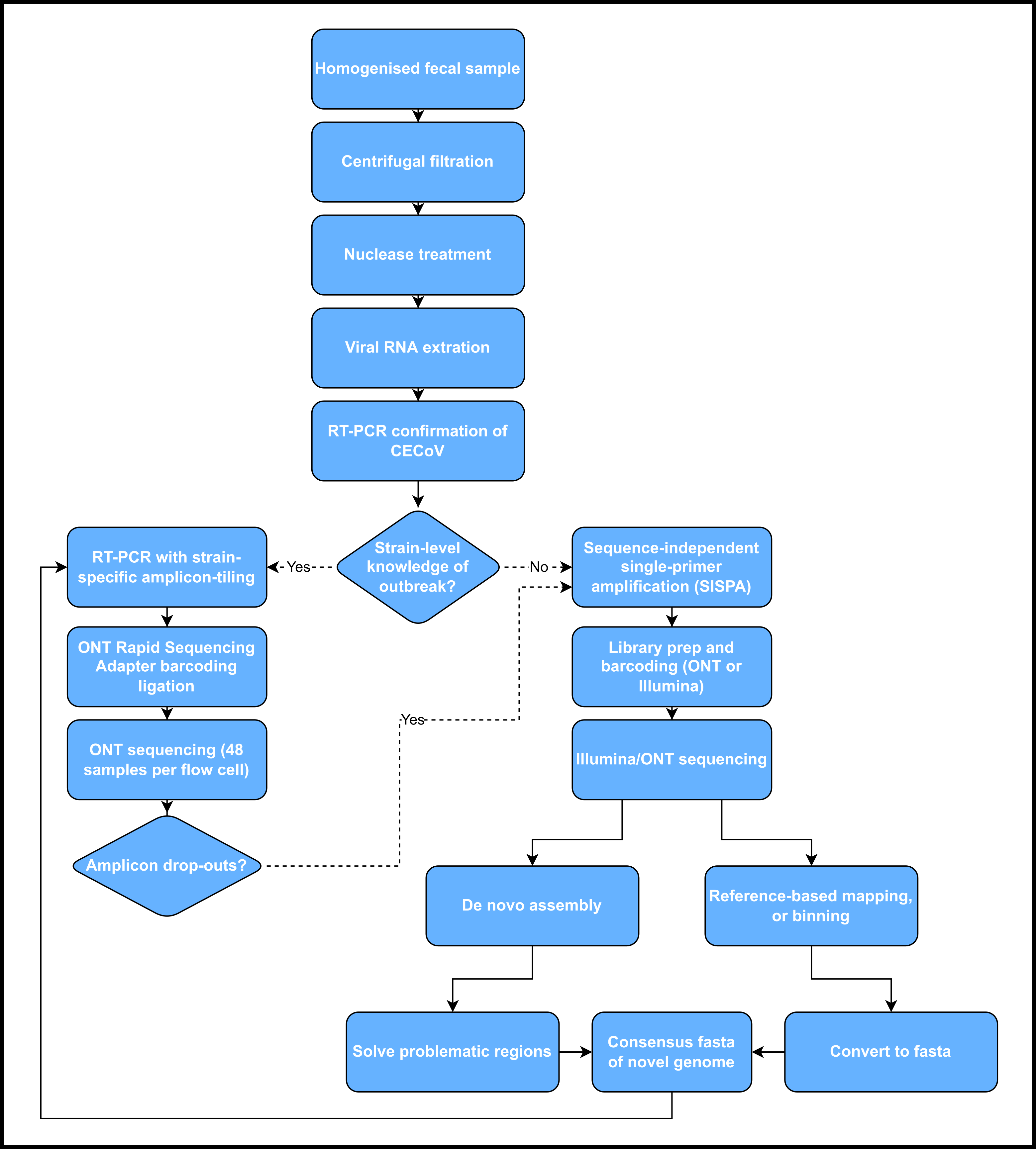
